# Genetic Background and Sex Modulate Androgen Responses in Human Brain Microphysiological System

**DOI:** 10.64898/2025.12.03.692130

**Authors:** Maren Schenke, Jason Laird, Alex Rittenhouse, Viktoriya Kucheryavenko, Winfried Neuhaus, Ou Chen, Sarven Sabunciyan, Alexandra Maertens, Lena Smirnova

## Abstract

Sex steroids shape human brain development, yet the cellular and molecular consequences of androgen addition need further exploration. Here, we used a human neural organoid model - brain microphysiological system (bMPS), derived from nine induced pluripotent stem cell (iPSC) lines to model the impact of dihydrotestosterone (DHT), a potent non-aromatizable androgen. All lines differentiated reproducibly into electrically active, neuron–glia–oligodendrocyte organoids expressing robust androgen-signaling components. DHT was bioavailable, elicited nuclear androgen receptor (AR) translocation and increased organoid size in most cell lines, consistent with androgen-responsive mTOR and metabolic pathway activation.

Bulk RNA-seq across nine lines revealed that transcriptional responses varied across donor backgrounds, but DHT-responsive genes converged on mitochondrial energetics, lysosomal function, glycoprotein processing, apoptosis, and mTOR signaling. Cell-type expression profiling showed an androgen-driven shift primarily in male lines toward astrocytic profiles with reductions in oligodendrocyte, oligodendrocyte progenitor (OPC), excitatory, and inhibitory neuronal signatures, supported by immunohistochemistry and AR enrichment in astrocytes and OPCs. DHT also altered neurodevelopmental pathways, increasing variation in synaptic pruning and decreasing variation in neuronal migration, with autism spectrum disorder (ASD) diagnosis and seizure status of donors moderating these effects more strongly than sex.

Baseline transcriptional differences distinguished iPSC lines which responded more strongly to DHT from weak responders: responders displayed enhanced synaptic maturity and reduced ECM gene expression. Using isogenic XX/XY lines, we found that differences in sex-chromosome expression exceeded DHT-induced changes and that DHT decreased expression of inhibitory neuron genes in males and increased it in females. Finally, DHT induced extensive DNA methylation changes, targeting *HOX* genes, patterning, and synaptic genes.

Collectively, these findings reveal that androgen signaling shapes transcription, cell populations, and epigenetic landscapes in a genetic background-dependent manner. This work contributes to understanding how androgens influence human brain development and highlights how *in vitro* models can contribute to representing inter-individual variability in neurodevelopment and neurodevelopmental disorders.

## Introduction

Sex as a biological variable influences human physiology, but a historic focus on male subjects in clinical trials or lack of consideration of sex as a variable (Geller et al., 2018) led to a significant knowledge gap. Despite a recent push to change this, sex differences are not fully covered or addressed in pre-clinical studies and especially in cell-based approaches (Beery & Zucker, 2011; Kouthouridis et al., 2022; Woitowich et al., 2020). Beyond sex chromosomes, sex hormones also influence human physiology as exemplified by a spike in testosterone in males, which coincides with a critical window of neurogenesis, influencing the development of the brain (van Hemmen et al., 2016, Brown, 2023). This early androgen exposure has long been hypothesized to modulate and shape long-term brain organization, and alterations in hormone exposure may contribute to sex-biased vulnerability to neurodevelopmental disorders (NDDs) (Baron-Cohen, 2002). The prevalence of NDDs such as autism spectrum disorder (ASD) have been on the rise for decades, but their etiology is not well understood (Hertz-Picciotto et al., 2018; Solmi et al., 2022). Recent studies found that one in 31 children in the US is now diagnosed with ASD (Maenner et al., 2020, 2023), with boys diagnosed about 4 times more than girls. While differences in symptom presentation, diagnostic criteria, and awareness contribute to this imbalance (Bölte et al., 2023; Loomes et al., 2017; Werling & Geschwind, 2013) it has also been speculated that biological sex and early hormone signaling influence developmental trajectories of neurodevelopmental disorders diagnosed during childhood. In mammals, who typically either have a XX or XY genotype, it is the SRY locus (sex-determining region of the Y chromosome), which initiates the formation of testes which results in the production and secretion of testosterone. Two major peaks of testosterone production occur in mammals: In pre- and perinatal development and then again with the onset of puberty. The first phase induces so-called long-term changes (organizational phase) that will become relevant with the second surge of testosterone (activational phase). As NDD diagnosis typically occurs during the first 4-5 years of life, it is likely that the sex bias in prevalence has its roots in the first testosterone surge (McCarthy et al., 2017). It is important to note that the absence of sex hormones during development results in formation of a female phenotype as a default (McCarthy, 2024). In rodents, the most commonly studied model, the sexual differentiation of the brain involves local aromatization of testosterone into estradiol, which is the main inducer of brain masculinization using downstream signaling mainly via the Estrogen Receptor alpha (ERalpha) (McCarthy, 2008; Zuloaga et al., 2008). Sex differences in neurophysiology and behavior in rodents are by now well studied which was eased by the use of the 4-core-genotype model, which separated the development of testis from the presence of the Y-chromosome by moving the SRY-gene to an autosome, allowing hormonal and chromosomal effects to be studied separately (De Vries et al., 2002). Most of the sex differences in rodent brains have been shown to be sex hormone induced and include changes in, for example, the size of specific brain areas, the extent of their connectivity, the number of synapses or (epi)genetic changes, that strongly influences the animals reproductive behavior (McCarthy, 2024). In humans and non-human primates however, this process shows a significant species-difference, where aromatase is less expressed and testosterone exerts its effects primarily through direct binding to the androgen receptor (AR) (McCarthy, 2024). Despite these species differences, most of our current knowledge in developmental biology is derived from animal models. In humans, sex differences of brain development are widely debated and appear to be less pronounced than in rodents (Eliot et al., 2021). Still, naturally occurring mutations have shown that the involvement of androgens and the androgen receptor are essential for the development of a masculinized phenotype, which has shown to extend to the brain. Nonetheless, the limitation of animal models, especially with regard to representing the human brain, leave major gaps in understanding of the molecular consequences of androgen signaling in the developing brain (van Hemmen et al., 2016, Bakker, 2022).

Human stem cell-derived models of the brain have been used to study NDDs successfully and were able to provide spatiotemporally resolution of neurodevelopmental trajectories on the molecular and cellular level (Birtele et al., 2025). While *in vitro* models cannot reproduce e.g. behavioral traits as found in ASD, they can help gain knowledge on the underlying neurophysiological aspects of neurological disorders (Kilpatrick et al., 2023). Recent work has highlighted that most human *in vitro* neural models do not address the impact of sex differences, despite sex being a variable in neurodevelopment: cell lines often lack reliable sex-chromosome integrity, hormonal milieus are rarely developmentally appropriate, and brain organoids typically are not designed or powered to interrogate sex-specific mechanisms (Castro-Aldrete et al., 2025). As a result, current *in vitro* systems only partially capture sex differences in neural development and disease vulnerability. This gap underscores the need to optimize human neural organoid models with controlled androgen exposure and well-annotated donor backgrounds, to mechanistically dissect how sex hormones and sex chromosomes shape human neurodevelopment. Here, we addressed the significance of androgen signaling in neurogenesis and neural development. To do so, we optimized and further characterized iPSC-derived neural organoids or brain microphysiological system (bMPS) by incorporating a potent, endogenous androgen, dihydrotestosterone (DHT) as a physiological component. Previously, this bMPS model was used to study diseases and neurotoxicants (Bullen et al., 2020; Modafferi et al., 2021; Morales Pantoja et al., 2024; Pamies et al., 2017, 2018, 2022; Romero et al., 2022). By treating differentiating neural organoids with DHT, we elucidate androgen signaling-specific molecular and cellular changes associated with brain development. For this, we utilize a set of nine cell lines: from male and female donors, from typically developed individuals, or from donors carrying an ASD risk mutation. Two of the cell lines are isogenic, and were derived from an individual with a mosaic form of Klinefelter’s syndrome (47,XXY/46,XY/46,XX), meaning that a line with XX and one line with XY genotype have been derived from the same individual (Waldhorn et al., 2022). Because androgen signaling can alter gene expression, cellular composition, and pathway activity, we applied a multi-omics strategy to resolve how it reshapes neurodevelopment across transcriptional programs, methylation states, cell populations in the context of different donor backgrounds.

## Results

### Functional Human bMPS reveal active androgen signaling with DHT-dependent effects on growth and Ca²⁺ dynamics

To model the effect of testosterone on human brain development, a total of 9 induced pluripotent stem cell (iPSC) lines from 8 different donors (Fig 1a and Supplementary Table 1) were differentiated into a brain microphysiological system (bMPS). Cell lines M1 and F1 were routinely used for this protocol, underwent quality control previously (Romero et al., 2022) and were used to establish the protocol used in this project.

**Fig. 1:**
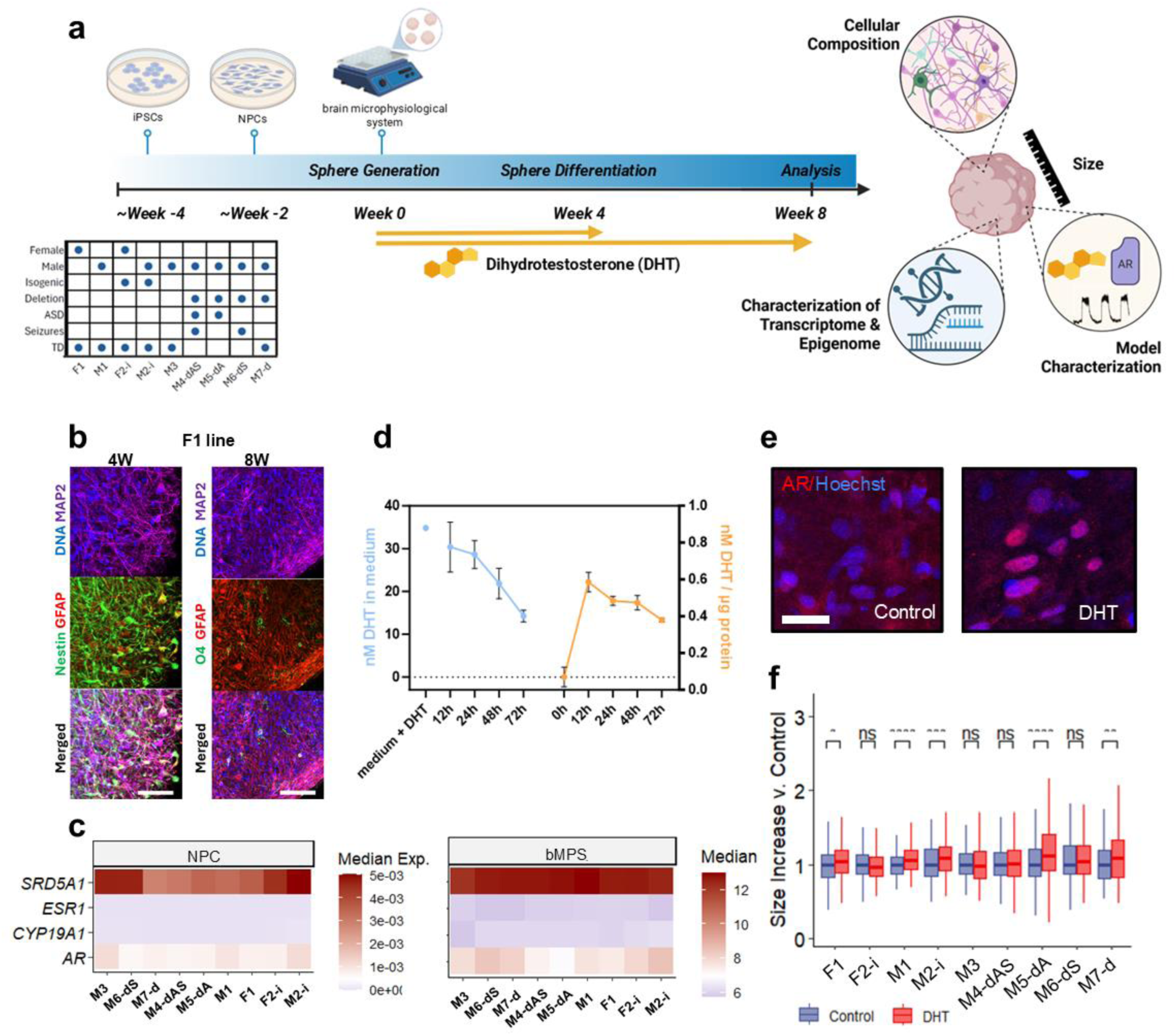
bMPS differentiation and characterization. **a)** Experimental overview detailing demographic and genetic background information on each cell line, brain organoid generation from iPSCs and DHT treatment timelines and analysis. **b)** Representative immunostaining of bMPS from the F1 iPSC line (other lines are shown in Fig. S1b), at 4W of differentiation and 8W of differentiation. Immature bMPS (4W) are positive for Nestin (neural stem cell marker), GFAP (glia marker), and MAP2 (neuronal marker), while mature bMPS (8W) are positive for MAP2, GFAP and O4 (oligodendrocyte precursor cell marker). Scale bar is 50 and 100 µM respectively **c)** Quantification of androgen-associated genes (*AR*, *SRD5A1*) and estrogen-associated genes (*ESR1*, *CYP19A1*) expression levels in neural progenitor cells by RT-qPCR and RNAseq at the 8W timepoint. **d)** Concentration of DHT in medium (blue) and in cell culture supernatant (orange). **e)** Immunostaining of androgen receptor (AR) in bMPS exposed to DHT or to solvent. Scale bar = 25 µm. **f)** Relative organoid size increases from day two to eight weeks comparing DHT-exposed organoids to solvent controls.

The remaining cell lines underwent the same steps of quality control. iPSC were karyotyped, were mycoplasma free, exhibited typical colony morphology and were positive for pluripotency markers. Efficiency of neural induction from iPSC to neural progenitor cells (NPC) was verified by quantification of Nestin- and SOX1-positive cells. Over 80% of the NPC were double positive across the lines, except M2-i with 79.1% and M3 69%, respectively (Fig. S1a). NPCs were differentiated into bMPS as initially described by Romero et al., (2023). To verify that the cellular composition and differentiation timelines were comparable, bMPS were immunostained at an intermediate timepoint of four weeks and at the end of the differentiation, at eight weeks (Fig 1b and S1b, c). All lines expressed neuronal marker MAP2, glia marker GFAP and neural progenitor marker Nestin at four weeks. At eight weeks of differentiation, all lines were positive for MAP2, GFAP and O4, a marker for oligodendrocytes with significant increase in the levels of MAP2 and GFAP, indicating appropriate maturation of the bMPS. Finally, electrical activity of the bMPS was assessed by measuring Ca^2+^ flux around week 8 of differentiation. Although patterns of the spiking were different between the cell lines, all cell lines demonstrated spontaneous electrical activity (Figure S1d).

Before addressing the effects of testosterone on neural differentiation, we first assessed the presence of receptors and enzymes necessary for testosterone metabolism and activity. In neural progenitor cells, expression levels of androgen receptor (*AR*), estrogen receptor alpha (*ESR1*), and enzymes involved in synthesis of DHT from testosterone (steroid 5α-reductase 1, *SRD5A1*) and of estrogens from androgens (aromatase, *CYP19A1*) were quantified via RT-qPCR. Expression levels showed the same trend at week 8 of the differentiation (Fig 1c). At both timepoints, expression levels of estrogen receptor and aromatase were several orders of magnitude lower than androgen receptor or *SRD5A1*, indicating a higher relevance of androgen-related signaling compared to estrogen-related signaling in our neural model. Supplementation of DHT, a testosterone metabolite which is a potent endogenous and non-aromatizable androgen, was initiated two days after seeding of NPCs for differentiation into bMPS, and organoids were treated with DHT either for 4 or 8 weeks. To assess whether DHT can be taken up by the cells, we quantified DHT concentrations via DHT-ELISA assay in both cell culture medium and bMPS lysates over the course of three days, from day 2 to 5 of bMPS differentiation (Fig 1d). From an initial DHT concentration of 60 nM, we were able to detect 35 nM of bioavailable DHT in fresh medium, which reduced over the time of exposure to 15 nM. We detected DHT in the lysates of washed bMPS, indicating uptake. This was corroborated by immunostaining for AR, which was, as expected, relocated to the nuclei of DHT-treated bMPS and not in the vehicle control cells (Fig 1e). Given that testosterone had been shown to influence proliferation (Quartier et al., 2018, Kelava et al., 2022), we measured organoid size as a readout of proliferative activity. Presence of DHT in culture medium significantly increased size of organoids in 5 out of 9 cell lines. The increase was slightly higher and more significant in M1 vs. F1 and was significant only in male isogenic line (M2-i) and not in the female isogenic line (F2-i) (Fig. 1f).

### DHT elicits heterogeneous transcriptional responses across donor backgrounds but converges on core cellular pathways

To investigate DHT-induced molecular changes, bMPS differentiated for eight weeks in presence of DHT or the solvent control were submitted to bulk RNA-seq, after verifying electrical activity via calcium imaging (Fig S1d). First, two treatment scenarios were assessed to determine the critical window of DHT effects. We treated cell lines F1 and M1 with 60 nM DHT either for four (week 1 to 4) or eight (week 1 to 8) weeks. Samples were collected at week 8 for both treatment groups. DMSO was used as a solvent control. We selected the eight week treatment as our standard treatment scheme for all subsequent experiments in the remaining seven iPSC lines. We analyzed the transcriptome and changes in DNA methylation in all cell lines differentiated under DHT supplementation or control samples as highlighted in figure 2a. DHT-induced transcriptional changes within a cell line were determined using differential expression analysis with *DESeq2* (Love et al., 2014). Genes with nominally significant changes (P<0.05, |FC|>1.5) were interrogated further (Supplementary Data 1). Interestingly, after 8 weeks of treatment with DHT, M1, M7-d, and M6-dS had more differentially expressed genes (DEGs) compared to the other cell lines (Supplementary Fig. S2a). Since several cell lines belonged to similar metadata categories (e.g. male or female), we quantified concordant DHT-induced transcriptional changes while accounting for line-specific variation using a random effects model. Using this approach, we identified several sets of DEGs per category (P < 0.05), with female and seizure sets having the greatest absolute number of DEGs and typically developing (TD) and ASD sets having the least (Fig. 2b; Supplementary Data 2). Two male cell lines, M5-dA and M3, did not have a strong transcriptional response to DHT, which may explain the lower number of DEGs in the TD and ASD sets (Supplementary Fig. S2a). Male, seizure, and 16p11.2 deletion sets had the most overlap between DEGs, likely driven by shared samples among these categories (Fig. 2b). Comparisons between individual cell line DEGs and random effects model DEGs found that the M1, M4-dAS, M7-d and M6-dS overlapped most with random effects DEGs, although this overlap was minimal (Jaccard coefficient < 0.05) (Supplementary Fig. S2b).

**Fig. 2:**
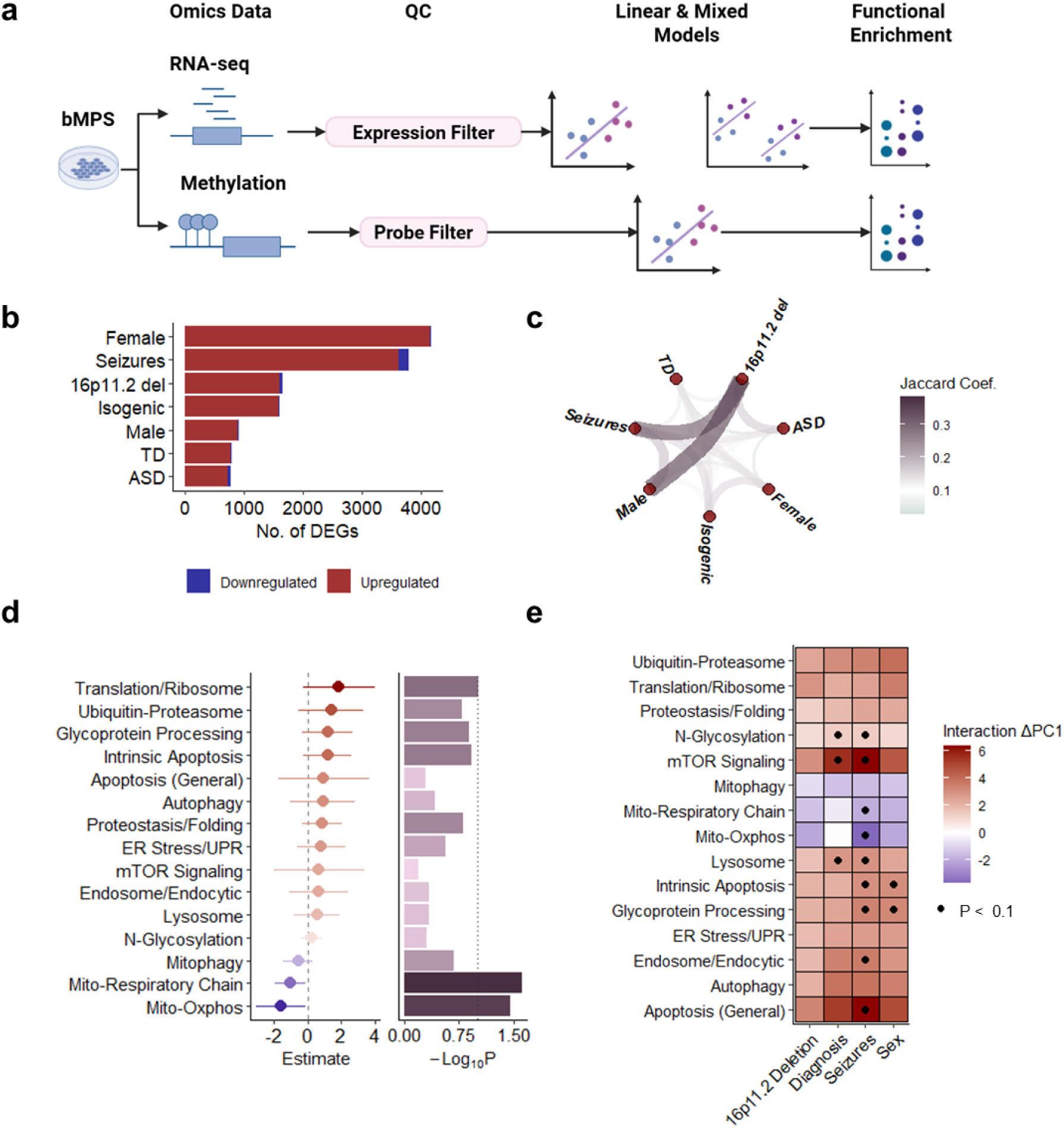
Global transcriptional responses to DHT. **a)** Data analysis pipeline describing multi-omics data pre-processing, modeling approaches and biological interpretation. Created in BioRender. Laird, J. (2025) https://www.biorender.com/. **b)** Number of DEGs identified using a random effects model (P < 0.1) stratified by demographic and genetic background. **c)** Pairwise overlap between random effects DEG sets colored by the Jaccard coefficient. **d)** Random effects model estimates (bottom) for DHT-associated biological processes colored by the change direction. The significance of each model (top) is represented by a barplot of the −𝐿𝑜𝑔(𝑃) values. **e)** Interaction term estimates between biological process PC1 and DHT exposure status, colored by change direction, and significant interactions (P <0.1) indicated by dots.

Each set underwent gene ontology (GO) enrichment analysis and significant terms (FDR < 0.1; Supplementary Data 3) were subsequently grouped into broader biological categories to improve interpretability (Gierlinski, 2025; Sayols, 2023).

Examination of overlapping GO categories revealed shared enrichment in mitochondrial energy metabolism, lysosomal gene expression, and mTOR signaling (Supplementary Fig. S2c). Increases in mTOR signaling, integral to growth and nutrient sensing, is consistent with the observation that organoids from most lines increased in size (Fig. 1f). While these sets did not overlap greatly at the gene level, transcriptional changes appear to have converged on common biological pathways. We interrogated these pathways further by examining the relationship between biological pathway variation and DHT exposure. DHT appeared to reduce variation in mitochondrial energetic pathways, which increased variation in glycosylation, lysosomal mechanisms, mTOR signaling, cellular stress responses, apoptosis, and transcriptional/translational programs (Fig. 2d). To understand if these relationships were modified by factors like sex, seizure status, ASD diagnosis, or 16p11.2 deletion status, we performed interaction analyses, setting the cell line as the random effect (Fig. 2e). In male lines, variation in glycoprotein processing and intrinsic apoptosis was increased compared to females. Seizure diagnosis was linked to increased variation in several common DHT biological processes, including apoptosis, endosomal gene expression, N-glycosylation, and mTOR signaling and decreased variation in mitochondrial energetics. ASD diagnosis was associated with increased variation in lysosome mechanics, mTOR signaling, and N-glycosylation. These results suggest that DHT-responsive biological processes vary across sex, ASD diagnosis, and seizure status, with the strongest pathway changes associated with seizures, followed by ASD diagnosis and sex (Fig. 2e). In contrast, the presence of the 16p11.2 deletion did not show a significant effect on the response to DHT (Fig. 2e).

### DHT reshapes cell composition and modulates neurodevelopmental processes depending on sex, ASD, and seizure status

Because DHT has been shown to influence neural lineage specification through androgenreceptor–dependent pathways (Kelava et al., 2022; Quartier et al., 2018), we next assessed whether DHT exposure altered cell-type composition within the bMPS. First, we fitted a linear mixed model to predict cell-type signatures, derived from the median expression of genes enriched in central nervous system (CNS) cell types, and set the cell line as the random effect. We found that the astrocyte profile increased with DHT treatment, while oligodendrocyte, oligodendrocyte precursor cell (OPC), excitatory and inhibitory neuronal signatures decreased (Fig. 3a). We confirmed RNA-seq findings with immunohistochemistry. RNA-seq analysis was in line with increase of GFAP and decrease of O4 signals (Fig. 3b and c). We then determined if DHT-induced cell-type changes depend on factors like sex, diagnosis of ASD, seizures, or 16p11.2 deletion status.

**Fig. 3:**
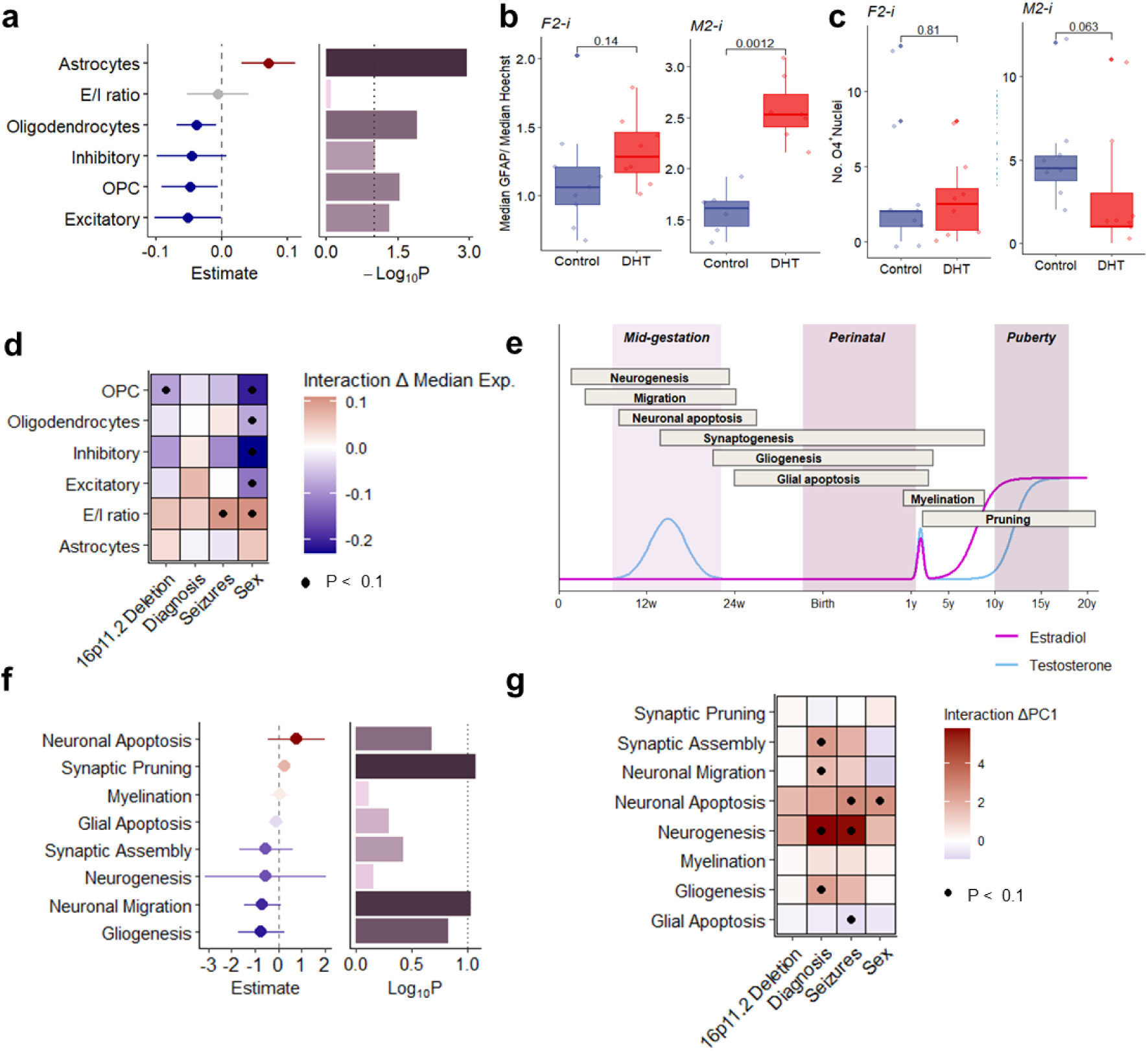
Impact of DHT treatment on cell marker and developmental set expression. **a)** Linear mixed model estimates (left) of DHT-induced shifts in CNS cell type signatures, colored by the change direction. The significance of these relationships is represented by a barplot of the −𝐿𝑜𝑔(𝑝 − 𝑣𝑎𝑙𝑢𝑒). **b)** Heatmap of the interaction term estimate with the cell type signatures, colored by change direction, and significant interactions (P <0.1) are labeled with dots. **c)** Change in GFAP immunofluorescence normalized to Hoechst intensity in the F2-i and M2-i lines between DMSO and DHT conditions. **d)** Number of O4-positive oligodendrocytes per organoid section in the F2-i and M2-i lines between DMSO and DHT conditions. **e)** Timeline of cellular neurodevelopmental processes and testosterone and estradiol peaks during development (Adapted from (Brown, 2023)). **f)** Linear mixed model estimates (left) of DHT-induced shifts in neurodevelopmental processes, colored by the change direction. The significance of these relationships is represented by a barplot of the −𝐿𝑜𝑔(𝑝 − 𝑣𝑎𝑙𝑢𝑒). **g)** Heatmap of the interaction term estimate with the cell type signatures, colored by change direction, and significant interactions (P <0.1) are labeled with dots.

DHT treatment led to downregulation of OPC-specific genes in cell lines with the 16p11.2 deletion and increase of the excitatory/inhibitory (E/I) ratio in cell lines derived from patients with seizures (Fig. 3d). The effects of DHT vary greatly by sex, where OPC, oligodendrocyte, inhibitory and excitatory profiles decrease, and the E/I ratio increases in males. Interestingly, the strongest change was the reduction in gene expression of inhibitory neuron and OPC signatures in male lines. As discussed above, neurodevelopmental processes coincide with elevated levels of hormones before and after birth (Fig. 3e). Using a linear mixed modeling approach, and setting the cell line as the random effect, we determined how DHT may affect neurodevelopmental processes. Based on transcriptomics data, we observed that DHT treatment increased variation in synaptic pruning and decreased variation in neuronal migration transcriptional programs (Fig. 3f). We next tested whether the relationship between DHT exposure and neurodevelopmental transcriptional variation is modified by sex, ASD, seizure and 16p11.2 deletion status. In contrast to cell marker profiles, the relationship between DHT and neurodevelopmental processes was predominantly moderated by seizure and ASD status (Fig. 3g), consistent with the GO findings (Fig. 2e). DHT increased variation in neurogenesis-related expression in lines derived from donors exhibiting seizures or ASD. Lines obtained from ASD-exhibiting donors amplified DHT-associated variation in synaptic assembly, neuronal migration, and gliogenesis. Seizure status was associated with increases in neurogenesis, neuronal apoptosis and reduced glial apoptosis in response to DHT. Male donor-derived lines showed greater DHT-associated variation in neuronal apoptosis. Collectively, these results show that the impact of DHT on neurodevelopmental transcription is not uniform but is rather shaped by ASD and seizure diagnoses of cell line donors, similar to the GO pathways enrichment results.

### Intrinsic transcriptional states underlie cell-line-specific sensitivity to DHT

The M1, M7-d, and M6-dS cell lines had the most DEGs in response to DHT compared to other cell lines. To better understand why some lines responded to DHT, we examined baseline (transcriptional profiles in vehicle control samples) across two groups of cell lines: the lines with a robust DHT response (Responder: > 200 DEGs) and those with reduced response (Nonresponder: < 200 DEGs). We compared the baseline expression profiles of each DHT-responder to all other non-responder lines, identified nominally significant DEGs (P < 0.05, |FC| > 1.5), and then kept only genes that changed in the same direction across all responder-non-responder comparisons (Fig. 4a; Supplementary Data 4). GO enrichment revealed that extracellular matrix related genes were largely downregulated, while synaptic gene expression was upregulated in responder cell lines (Fig. 4b; Supplementary Data 5). This prompted us to interrogate focused gene sets. We found that responders had greater expression of genes involved in androgen biosynthesis and synaptic genes (both glutamatergic and GABAergic), while expression of genes related to extracellular matrix (ECM) was reduced (Fig. 4c). Greater expression of synaptic genes suggested that DHT-responders may be more transcriptionally mature at baseline.

**Fig. 4:**
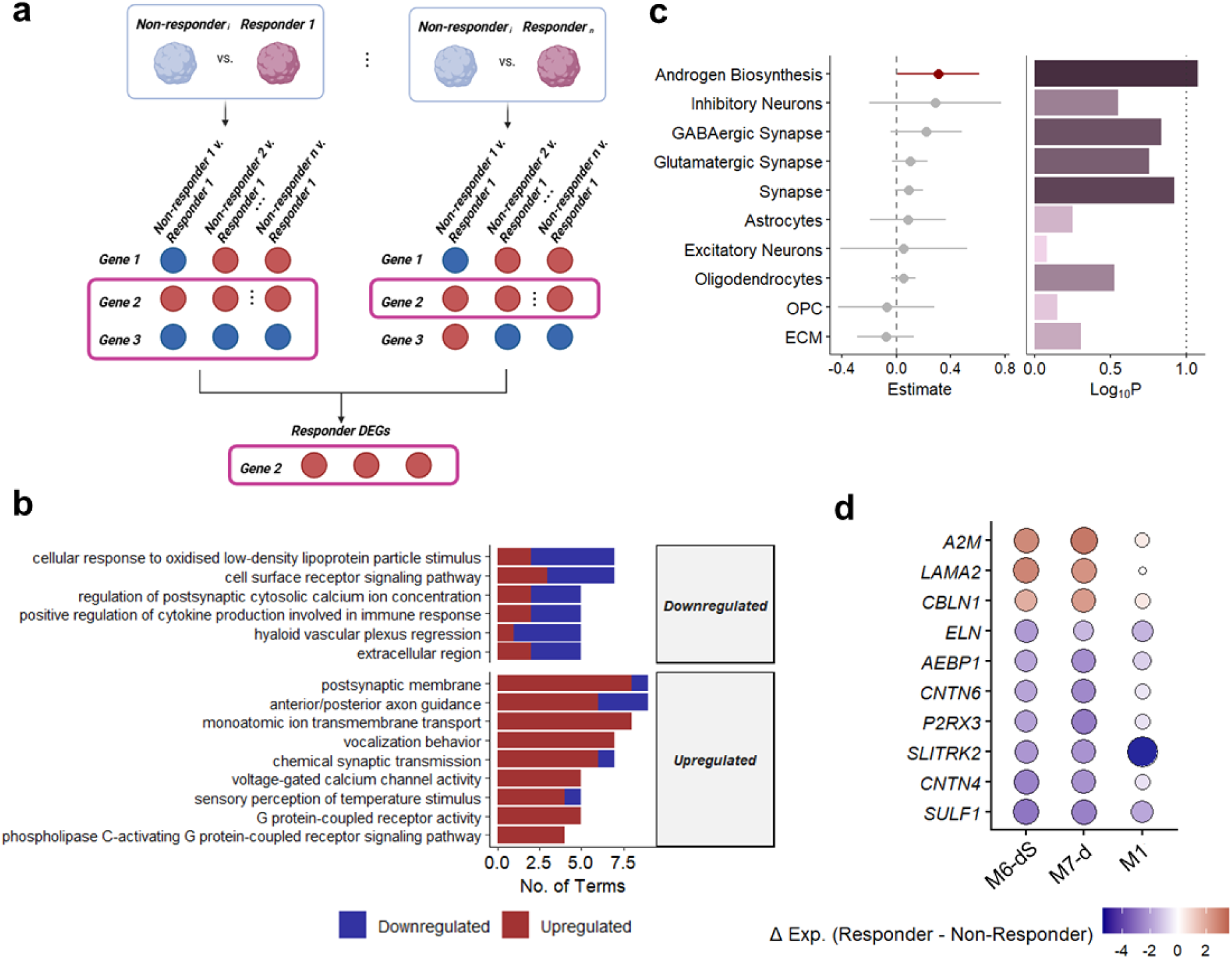
Baseline transcriptional differences between DHT-responders and non-responders. **a)** Identification of DHT-responder DEGs, by performing pairwise differential expression between each responder with every non-responder, then filtering DEGs by genes with consistent change directions within responder-non-responder comparisons. Created in BioRender. Laird, J. (2025) https://www.biorender.com/. **b)** GO enrichment of DHT-responder DEGs, colored by change direction showing an upregulation of genes related to synaptic gene expression and reduced ECM gene expression in DHT responders. **c)** Linear mixed model effect estimates for DHT-responder associated changes in biological processes associated with DHT responders showing elevated androgen biosynthesis and synaptic gene expression in DHT responders. **d)** Top DHT responder DEGs colored by change direction.

Examination of the genes with the greatest fold change between non-responders and responders revealed that *A2M*, *LAMA2*, and *CBLN1* were upregulated and *ELN*, *AEBP1*, *CNTN6*, *P2RX3*, *SLITRK2*, *CNTN4*, and *SULF1* were downregulated (Fig. 4d). These represent proteins responsible for synapse organization (*CBLN1*) and synaptic adhesion (*CNTN4*/*CNTN6*) (Mikulska-Ruminska et al., 2017; Yuzaki, 2011).

### Transcriptional influence of sex chromosomes vs DHT in genetically matched isogenic XX and XY iPSC lines

Since known iPSC-donor variability still poses a challenge, we used isogenic cell lines, generated by Waldhorn et al., (2022) from a patient with Klinefelter’s syndrome. These represent a great model to address the effects of DHT on neural development while accounting for genetic background. Additionally, using these lines afforded us the opportunity to disentangle transcriptional changes driven by sex chromosomes vs. DHT treatment while controlling for genotype (XX and XY). We first compared the baseline transcriptional differences between isogenic F2-i and M2-i lines (vehicle control) and then compared the number of DEGs to DHT-induced DEGs within cell lines. Sex-chromosome differences (XX v. XY) between the isogenic lines produced more DEGs than DHT exposure in the F2-i vs M2-i lines (Fig. 5a; Supplementary Data 6). GO enrichment analysis of chromosomal DEGs at the baseline (F2-i vs. M2-i) revealed broad downregulation of GABAergic gene expression and cell adhesion, whereas transcriptional machinery genes were upregulated in the M2-i line (Fig. 5b; Supplementary Data 7). Because GABAergic-specific processes were reduced in the M2-i line, we next examined the median expression of genes enriched in inhibitory neurons across all cell lines, both at baseline and after DHT treatment. We found that baseline expression of inhibitory neuron signatures was not consistently lower in male-derived lines (Fig. 5c). However, DHT decreased expression of inhibitory neuron genes in males and increased it in females, when compared to the baseline expression (Fig. 5c). When we interrogated which GABAergic genes drove this signal, we found that the effect stemmed from GABAergic receptor expression rather than GABAergic synapse components (Fig. 5d). Together, these findings highlight that both sex chromosomes and DHT exposure modulate transcription during neural differentiation, with the former having the greater effect.

**Fig. 5:**
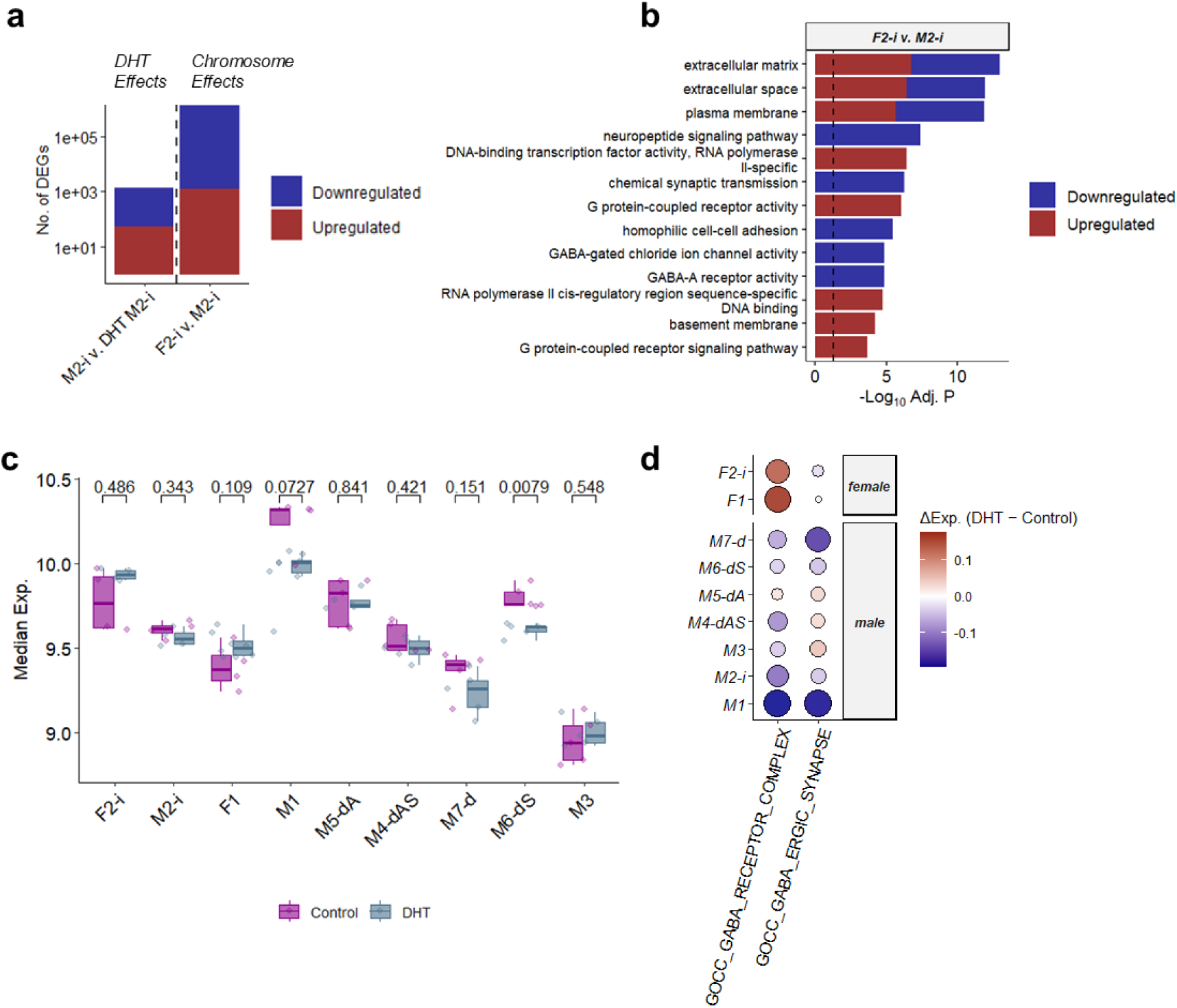
Sex chromosome and DHT transcriptomic effects. **a)** Comparison of the number of DEGs by sex chromosome (F2-i v. M2-i, first row) versus DHT treatment in the individual cell lines, showing a much stronger effect of sex chromosomes. **b)** Top GO enrichment terms associated with sex chromosome DEGs, as derived from the isogenic comparison, where change direction is relative to the female isogenic line. **c)** Inhibitory neuron cell profile expression, derived by taking the median of inhibitory neuron-enriched genes, across cell lines and DHT exposure status. **d)** Change in GABAergic cellular component median expression between DHT and DMSO conditions for each cell line.

### DHT triggers extensive epigenetic remodeling, targeting patterning and synaptic genes

Because transcriptional responses provided only a partial view of DHT-mediated effects, we next examined whether androgen exposure also reprograms the epigenetic landscape. To this end, we profiled DNA methylation across four cell lines (M1, F1, M2-i and F2-i) to determine the extent and specificity of DHT-induced epigenetic remodeling. We performed differential methylation analysis using *limma-trend* (Ritchie et al., 2015). Nominally significant probes (P < 0.05, |FC| > 1.5) were retained for further analysis (Supplementary Data 8). Similar to the transcriptional analysis, we first analyzed the methylation changes in two lines, M1 and F1 treated with DHT for either 4 or 8 weeks, with samples processed at week 8 for both treatment groups.

M2-i and F2-i lines were treated only for 8 weeks. Regardless of treatment window, M1 had the greatest number of differentially methylated probes (DMPs) followed by F1, F2-i, and M2-i (Fig. 6a). Overall, methylation changes were more extensive than gene expression changes, indicating broader epigenetic remodeling prior to a transcriptional response (Fig. 6b). We next examined which genes were differentially methylated. DHT exposure altered the methylation of several *HOX* genes (Fig. 6c), a family of transcription factors which regulate anterior-posterior patterning (Nelms & Labosky, 2010). Among these, *HOXA5* contained the most differentially methylated probes, and was hypermethylated (Fig. 6c-d).

**Fig. 6:**
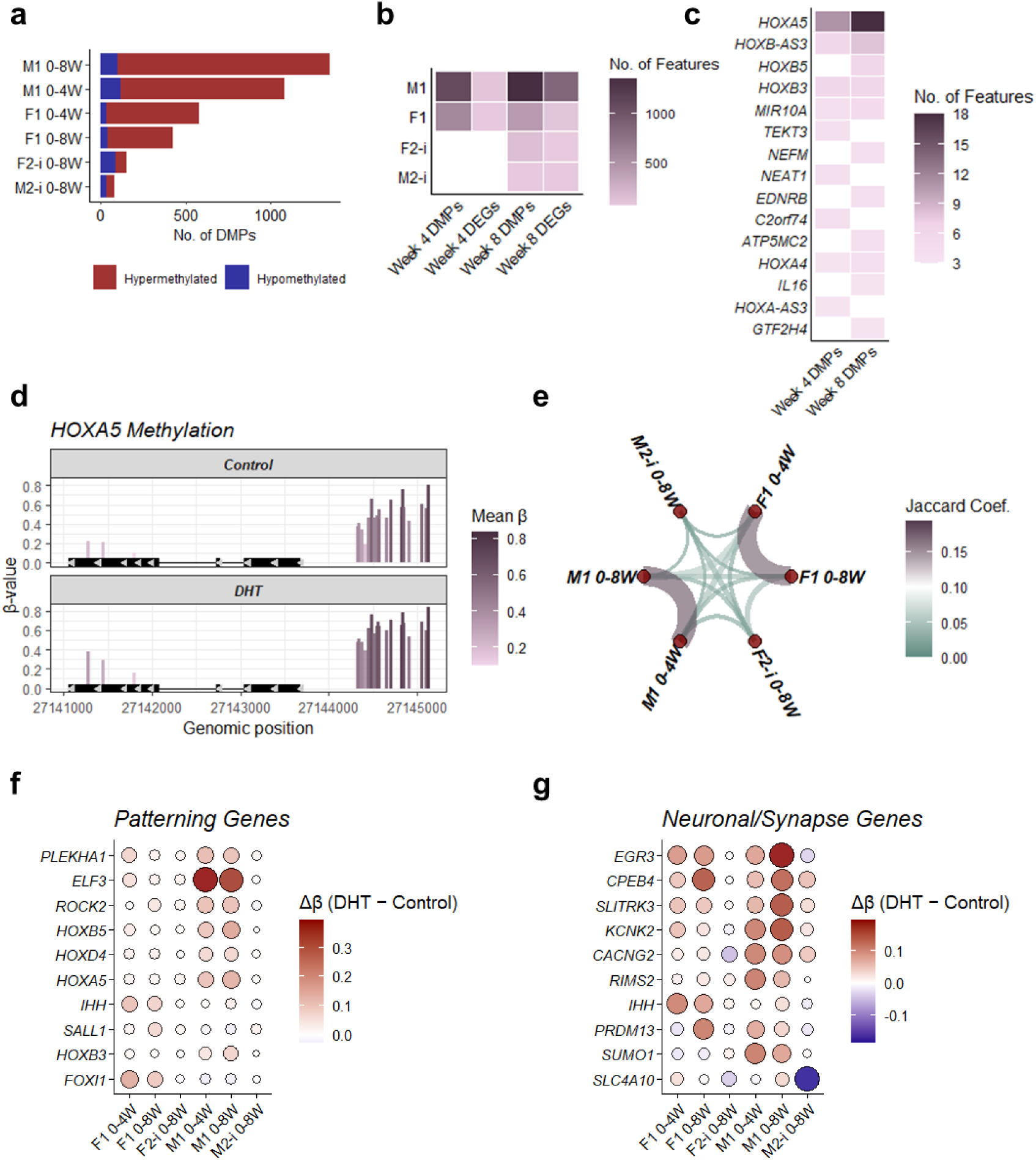
DHT-induced methylation changes across exposure timepoints. **a)** Number of differentially methylated probes per cell line and exposure time duration (4 and 8 week DHT exposure). **b)** Number of DHT-associated DMPs and DEGs across cell lines and exposure times. **c)** Frequently differentially methylated genes across exposure times. **d)** Representative change in methylation in *HOXA5* methylation in the M1 line (8 week DHT exposure) between DMSO and DHT conditions. **e)** Pairwise overlap in differentially methylated genes colored by the Jaccard coefficient. Top differentially methylated patterning genes **(f)** and neuronal or synapse-related genes **(g)** across cell lines and exposure times.

Overlap analysis revealed that differentially methylated gene overlap was highest within M1 and F1 cell lines (Fig. 6e). GO enrichment analysis of DMP-associated genes indicated that patterning, developmental processes, and neuronal/synapse genes were primarily hypermethylated (Supplementary Data 9), consistent with the transcriptomic evidence for synaptic remodeling. Several patterning genes with the strongest methylation changes, *PLEXHA1*, *ELF1*, *ROCK2*, *HOXB5*, *HOXD4*, *HOXA5* were hypermethylated in the M1, and *IHH*, *SALL1*, and *FOXI1* in F1 line (Fig. 6f). Interestingly, the extent of hypermethylation did not differ substantially between 4 and 8 week DHT exposures (Fig. 6f-g), suggesting that the first four weeks of differentiation are a critical developmental window for DHT treatment, which is sufficient to induce stable epigenetic reprogramming. In contrast, the F2-i and M2-i isogenic lines had a much weaker methylation response to DHT when compared to M1 and F1, which correlates with RNA-seq data and may reflect the clinical background of the donor, an individual with Klinefelter’s syndrome who was on testosterone replacement therapy. Across M1 and F1 lines, neuronal and synapse genes, like *EGR3*, *CPEB4*, *SLITRK3*, *KCNK2*, *CACNG2*, and *RIMS2*, were hypermethylated. Together, these findings indicate that DHT treatment, even during early stages of neurogenesis, induces significant changes to patterning and neuronal/synaptic genes.

## Discussion

Despite significant sex differences in brain disorders and diseases, there is a lack of consideration of sex as a biological variable in human *in vitro* models used to study them as standard culture methods do not recapitulate sex differences (Castro-Aldrete et al., 2025; Pavlinek et al., 2024). As MPS and other complex human-relevant models advance toward capturing inter-individual variability and personalized medicine, sex becomes an essential variable and needs to be explicitly incorporated within *in vitro* systems. Research has shown that sex hormones, especially testosterone, play a significant role in the development of the male human brain, even though the underlying molecular pathways are not entirely understood as data from animal experiments can not be fully translated to humans (van Hemmen et al., 2016, Bakker, 2022).

In this study we developed an *in vitro* model of the human brain, treated with DHT to model the potential masculinizing effects of physiological sex hormones. We used a diverse set of cell lines, demonstrating that not only the presence of the Y chromosome but also the presence of the androgen hormone, DHT, modulates neural differentiation *in vitro* through androgen receptor signaling. In humans, endogenous testosterone is reduced to DHT - an active non-aromatizable form with greater affinity to the androgen receptor. While testosterone, acting through androgen receptor signaling, is thought to be the predominant sex hormone during brain development in humans (Bakker, 2022), in rodents it is estradiol (aromatized from testosterone) and the estrogen signaling pathway (McCarthy, 2024). Here, we demonstrate high expression levels of androgen-signaling related genes with relatively muted expression of estrogen receptor and aromatase in neural organoids, showing that neural organoids capture this human-specific aspect of neural development. We verified that DHT is bioavailable in our culture system, is taken up by our model and then investigated the effect of DHT on neural differentiation in lines derived from female and male donors carrying diverse genetic backgrounds and relevant diagnoses. In particular, we assessed transcriptional, epigenetic and cellular responses across bMPS generated from nine different iPSC lines. Although individual cell lines varied in the magnitude of DHT-induced changes, shared biological processes converged on consistent themes. At the transcriptional level, DHT increases transcription, translation and mTOR signaling - core components of cellular growth. We found a small, but significant increase in organoid size in most DHT-exposed lines compared to the respective controls. This is in line with reported androgen-induced proliferation of neural stem cells and larger brain size in men (Eliot et al., 2021; La Rosa et al., 2021) and is consistent with previous work assessing androgen exposure in brain organoids in one male and one female line (Kelava et al., 2022). We find that effects of DHT on cell type composition are sex specific, while neurodevelopmental pathways are instead modified by seizure history, ASD diagnosis and 16p11.2 deletion status. We also found that the baseline transcriptional state predicted DHT responsiveness. Lines that mounted the strongest response, shared elevated androgen biosynthesis and higher synaptic gene expression, features consistent with greater neuronal maturity (Yuan et al., 2024). These findings suggest that the degree of maturation may prime cells to respond more to androgen exposure.

Using isogenic XX and XY lines, derived from an individual with a mosaic karyotype (Waldhorn et al., 2022), allowed us not only to more precisely control genetic background but also to separate the effects of sex chromosome and sex hormones. Comparing these two isogenic lines (XX vs. XY) derived from the same donor, we found that sex chromosomes exert a stronger effect on transcription than DHT treatment. However, since we use only a single donor for this comparison, and since we found that the cell lines used in this study show different magnitudes of DHT-induced changes, repeating this comparison in more lines would be necessary. The donor of the isogenic lines had Klinefelter syndrome and received testosterone therapy, both of which may have influenced our results. Combining all lines with a male genotype, we found reduced baseline GABAergic expression when DHT was supplemented. Postmortem transcriptomic studies have found that males have reduced expression of neuronal-associated genes and increased expression of astrocyte and microglial transcriptional programs (Werling et al., 2016). While our model does not contain microglia, we did find that DHT consistently upregulated astrocyte-related gene expression and increased GFAP-intensity in immunostaining of isogenic lines, especially in the male line, while oligodendrocytes appear to be downregulated. This supports the idea that DHT exposure may contribute to increased astrocyte populations in males. When examining neuronal expression trends, we found that DHT modulated inhibitory signaling in a sex-specific manner, by decreasing GABAergic receptor expression in most male lines while increasing it in female lines. This differential modulation suggests a mechanism through which androgen exposure may contribute to sex-biased neurodevelopmental phenotypes by altering E/I balance. In rodents, the prenatal surge of sex hormones has been shown to modulate GABAergic circuit function and GABAergic neurons contribute to the sexual differentiation of the nervous system, but specific effects depend on the specific brain area (McCarthy, 2024; Wolf et al., 2022). Sex-dependent and DHT-sensitive changes in inhibitory neuronal trajectories observed in our study may reflect the potential disbalance in E/I ratio and as a result male bias in ASD prevalence. The diagnosis-specific effects, particularly in ASD- and seizure-derived lines, underscore the potential relevance of androgen sensitivity to neurodevelopmental disorders with marked sex biases. Altered prenatal levels of sex steroids have been reported in subsets of ASD patients in epidemiological and clinical studies (Auyeung et al., 2009; Baron-Cohen et al., 2015, 2020), and hyperandrogenic conditions (e.g. maternal polycystic ovary syndrome PCOS or Congenital Adrenal Hyperplasia (CAH)) have been associated with increased ASD risk in some studies (Dooley et al., 2022; Dubey et al., 2021). Our finding that DHT disproportionately alters inhibitory signaling in male lines and increases neurogenesis-related variation in ASD- and seizure-derived lines corroborated findings that androgen signaling may interact with genetic liabilities to shape E/I balance, one of the best-established cellular signatures of ASD (Culotta & Penzes, 2020; Trakoshis et al., 2020).

At the epigenetic level, androgen signaling has been shown to modulate DNA methylation (Ammerpohl et al., 2013). In our study, DHT produced extensive and durable DNA methylation changes, exceeding the magnitude of transcriptional effects. Consistent with prior evidence that androgen-associated methylation changes can emerge long after exposure (Ghahramani et al., 2014), our data showed that both intermediate four-week DHT treatment during early neurogenesis phase (week 0 to 4) as well as longer exposure (week 0 to 8) induce methylation changes across patterning, developmental and neuronal gene programs. Intriguingly, *HOX* genes, key regulators of anterior-posterior identity, were among the most affected. This is particularly interesting as Gandal et al. found that in conditions like ASD, the anterior-posterior gradient is attenuated in the cerebral cortex (Gandal et al., 2022). This may raise the possibility that androgen exposure, or differences in androgen sensitivity may interact with genetic risk factors to alter regional identity.

Previous research on the effects of sex hormones and chromosome-dependent sex-differences could only provide partial answers to what extent these factors influence neural development. A study using four male and four female embryonic human embryonic stem cell lines differentiated into neurons found sex differences in gene expression, but did not supplement sex hormones (Pottmeier et al. 2024). A study by Bramble et al., (2016) found differences in expression levels of e.g. epigenetic regulators depending on sex chromosomes and androgen exposure, but used only murine cells. Other studies using immortalized neural cell lines or human neural stem cells found effect sizes to be small after short-term exposure to androgens, but describe effects on regulation of cell growth and cell differentiation (Quartier et al., 2018). A single study also used brain organoids as a more sophisticated model and treated it with sex hormones, and found a positive effect on cortical progenitor proliferation (Kelava et al. 2022). However, this study was conducted in one female iPSC line, with some endpoints measured in one additional male line. While our study significantly contributed to closing some gaps in our understanding of sex as a biological variable in brain development, it is not without limitations. First, the sample size within each genetic background/sex is relatively small, which reduces power to detect more subtle changes in transcription and methylation. Additionally, several of our cell lines have overlapping metadata categories, for example, all 16p11.2 deletion lines were derived from male donors. Our model does not include microglia, a cell type known to actively contribute to sex differences in mice, and to play a significant role in neurodevelopmental disorders such as ASD (Lenz et al., 2013; McCarthy, 2024; Velmeshev et al., 2019). This limits our ability to capture DHT-induced neuronal-immune interplay. Despite these limitations, our study also has several strengths. Using an *in vitro* model of the developing brain, we can model the effects of DHT under controlled conditions, excluding confounding effects of environmental and lifestyle factors. We examine DHT responsiveness across multiple cell lines representing diverse genomic backgrounds, allowing us to assess the robustness and heterogeneity of transcriptional and epigenetic effects. Integrational assessment of gene expression, DNA methylation and confirmation of transcriptional changes on the cellular and protein level provides complementary insights into DHT-induced regulatory changes in neural development. Incorporating random effects and interaction modeling enabled us to identify shared DHT effects and those that vary across genetic backgrounds and/or diagnoses. Together, these strengthen the interpretability and translational relevance of our findings.

Beyond examining the role of androgens in human neurodevelopment, our platform provides a foundation for testing how endogenous or exogenous hormonal environments (e.g. prenatal stress, endocrine disruptors, hormonal therapies) might shape early brain development. Such mechanistic insights are essential for interpreting sex-biased vulnerability to neurodevelopmental conditions or exposures and for developing personalized therapeutic strategies accounting for genetic background and hormonal milieu.

Our findings suggest that DHT shifts the intensity of core neurodevelopmental processes, such as synaptic maturation, metabolic state, and translational activity and drives broad, stable DNAmethylation changes. DHT is bioavailable, activates AR signaling, and shifts lineage balance toward astrocytes while reducing neuronal and oligodendrocyte programs, with stronger effects in males and ASD- or seizure-linked donors. The relationship between DHT and these shifts depends heavily on genetic background and sex. Using XX/XY isogenic lines demonstrated that sex-chromosome complement had a greater transcriptional impact than DHT treatment. Together, these results provide a more granular understanding of how DHT may influence the pace and coordination of neurodevelopment, and shape vulnerability to neurodevelopmental disorders. To our knowledge, this is the first study that systematically integrates androgen exposure, sex-chromosome complement, transcriptional programs, and DNA methylation across a genetically diverse set of cell lines in human brain organoids.

## METHODS

### Brain microphysiological system

All cell lines were cultured in an incubator on either Vitronectin- or Matrigel coated plates or flasks and checked for mycoplasma contamination regularly. For differentiation, either previously generated NPCs (M1 and F1) or iPSCs obtained from different repositories were used (See Supplemental Table 1). All lines were karyotyped and were mycoplasma free. iPSC lines were cultured in mTeSR Plus (StemCell Technologies) at 5% O_2_, 5% CO_2_, and 37°C. iPSCs were differentiated into NPCs in serum-free PSC Neural Induction Medium (Gibco, Thermo Fisher Scientific) following manufacturer recommendations and previously described protocol by Romero et al. (2022) and cultured at 37°C, 5% CO_2_ and 95% rel. humidity. NPC status was verified with flow cytometry for SOX1 and Nestin (see Fig. S1). For bMPS differentiation, NPCs were dissociated and plated in Neural Induction Medium at 2·10^6^ cells per well in 6-well plates under constant gyratory shaking (88 rpm, 19 mm orbit) to form aggregates. After 48 h, the medium was changed to differentiation medium (B-27 Plus Neuronal Culture System, 1% Glutamax (Gibco, Thermo Fisher Scientific), 0.01 μg/ml human recombinant GDNF (Gemini Bio), 0.01 μg/ml human recombinant BDNF (Gemini Bio), 1% Pen/Strep/Glu (Gibco, Thermo Fisher Scientific)). bMPS from each well were split into two groups: one was exposed to 60 nM DHT (Sigma, use of the controlled substance was approved under the laboratory’s DEA registration and supervised according to institutional controlled-substance handling procedures); the other group was used as a vehicle control with 0,0005% DMSO. About 75% of the medium was exchanged three times a week until the bMPS differentiation was concluded at week 8. DHT was added at each medium change, unless specified otherwise in the description of the experiment. Treatment continued for either 4 or 8 weeks.

### Flow cytometry

For characterization via flow cytometry, adherent cells were dissociated into a single-cell solution. Cells were fixed with 4% Paraformaldehyde (PFA) for 10 min at 4°C, followed by permeabilization for 10 min with 0,5% Triton X-100. Cells were washed, blocked and stained for 1 h with conjugated antibodies (See Supplementary Table 2). A LSRII flow cytometer (BD Biosciences) was used for sample analysis. Single-color and unstained samples were used as controls for compensation and gating.

### Immunostaining and confocal imaging

bMPS were fixed for 1 h in 4% PFA in PBS at 4°C. For staining, bMPS were permeabilized with 0.1% Triton in PBS for 30 min. then blocked with BlockAid™ Blocking Solution (ThermoFisher Scientific) for 1 h at 4°C. Staining with primary antibodies (See Supplementary Table 2) was performed overnight at 4°C and antibodies were diluted in PBS with 0.1% Triton, 0.5% BSA and 10% BlockAid. After three washes with PBS with 0.1% Triton, 0.5% BSA, bMPS were stained with secondary antibodies overnight at 4°C, washed three times and mounted with Immumount (Fisher Scientific). Nuclei were stained with Hoechst 33342 trihydrochloride (Invitrogen Molecular Probes) at a concentration of 1:10,000. Organoids were imaged on a Zeiss LSM800 confocal microscope.

### Quantification of DHT concentration from supernatants and lysates

DHT- levels in medium and cell culture supernatants were directly determined using a competitive ELISA (#IB59116, IBL America), while bMPS were first lysed via sonication and diluted lysates were included in the ELISA assay. Protein concentration of the lysates was determined via the Micro BCA™ Protein Assay Kit (Thermo Fisher Scientific) and extrapolated from a BSA standard dilution series. DHT concentration in lysates was normalized to the protein concentration as determined by the BSA assay.

### Calcium imaging

Neuronal activity was visualized using a calcium indicator. For this, bMPS were incubated with 10 µM Fluo-4AM (Tocris) and 0.5% Pluronic acid (Invitrogen) for two hours at 37°C. After washing, samples were transferred to glass-bottom plates and imaged on an Olympus FV3000-RS for 4-6 min at 12.5 frames per second. Raw fluorescence is depicted as ΔF/F.

### DNA and RNA extraction and library preparation

Adherent cells were washed with PBS and collected via scraping in lysis buffer on ice, while bMPS were collected, washed with ice-cold PBS, and pellets were flash frozen and stored at -80°C. Total RNA was extracted using the Quick-RNA^™^ Microprep Kit (Zymo Research) and DNA was extracted using the PureLink™ Genomic DNA Mini Kit (Thermo Fisher Scientific) according to the manufacturer’s instructions. Genomic DNA from bMPS differentiated for 8 weeks was diluted in the provided elution buffer and analyzed for DNA methylation using the EPIC BeadArray (Illumina). RNA from NPCs was used for RT-qPCR, while RNA from bMPS differentiated for 8 weeks was used to prepare libraries for Bulk RNA-seq as described previously (Alam El Din et al., 2025). Briefly, RNA concentration was quantified on a Qubit 3.0 and quality checked on the Agilent TapeStation 4200. For library prep, 500 ng of RNA were used with the TruSeq Stranded mRNA Library Prep Kit (Illumina), including poly-A selection, mRNA fragmentation, cDNA synthesis, 3’-end adenylation, and ligation of TruSeq RNA dual indexes. Libraries were amplified (15 cycles), purified, quantified on a Qubit 3.0, and assessed again on the TapeStation. After normalization and pooling, sequencing was performed on a NovaSeq X Plus (Illumina) using 150 bp paired-end reads. The sample preparation, DNA and RNA extractions and subsequent steps were performed in two batches, with analysis of M1 and F1 lines in the first batch, and all other lines in the second batch. The M1, F1, M2-i, F2-i lines had 4 replicates per cell line per DHT status, and all other lines had 5 replicates per cell line per DHT status.

### Reverse transcription and RT-qPCR

500 ng of total RNA were reverse-transcribed into cDNA in a volume of using M-MLV Reverse Transcriptase and Random Hexamer primers (Promega) according to the manufacturer’s instructions. For RT-qPCR, cDNA was amplified with 0.25 µM of forward and reverse primers (See Supplementary Table 3) and Fast SYBR™ Green Master Mix (Thermo Fisher Scientific) on a 7500 Fast Real-time PCR System to determine the quantification cycles (Cq). Relative gene expression levels were calculated with the 2^-dCT method (Schmittgen et al., 2008) and normalized to the geometric mean of the reference genes actin beta (ACTB) and cyclophilin A (PPIA). Suitable reference genes were selected using the geNorm algorithm implemented in the qBase+ software, version 3.4 (Biogazelle, Zwijnaarde, Belgium, www.qbaseplus.com) (Vandesompele et al., 2002)).

### RNA-seq processing

Raw sequencing reads were processed using the Nextflow nf-core/rnaseq workflow and containerized using conda (version 23.10.1) (Patel et al., 2025). Initial sequence quality control was performed with FastQC and Trim-Galore was used for removal of low-quality bases (*Babraham Bioinformatics - FastQC A Quality Control Tool for High Throughput Sequence Data*, n.d., *Babraham Bioinformatics - Trim Galore!*, n.d.). Reads were aligned with STAR to the human GRCh38 reference genome with Ensembl 111 release gene annotations (Dobin et al., 2013; Dyer et al., 2025). Post-alignment processing included coordinate sorting and indexing using SAMtools and PCR duplicate analysis using Picard (Li et al., 2009; Picard, n.d.). Gene and transcript quantification was performed with Salmon (Patro et al., 2017). Mapping and library quality control metrics were assessed using RSeQC for read distribution and insert size, Preseq for library complexity, dupRadar for duplication rates, BEDTools for coverage, and Qualimap for alignment quality (García-Alcalde et al., 2012; Quinlan & Hall, 2010; Sayols et al., 2016; *The Smith Lab*, n.d.; Wang et al., 2012). Gene counts were processed using the *tidybulk* R package, where genes with less than 1 count per million in 70% of the samples were filtered out for further processing (Mangiola et al., 2021).

### Methylation processing

DNA methylation was profiled using the Illumina Infinium Methylation EPIC v2.0 BeadChip array. Raw IDAT files were processed using the *minfi* R package (Aryee et al., 2014). Probes were filtered to remove any CpG site that failed in any sample based on detection p-values. Following initial probe filtering, samples were mapped to the human genome and normalized using preprocessFunnorm, producing normalized methylated and unmethylated signal intensities (Aryee et al., 2014). B-values were extracted and filtered to include probes with variance greater than 0.01. For statistical modeling, beta values were converted to M-values (logit-transformed).

### Identification of DHT-induced features

DHT-induced transcriptional and methylation features were determined by cell line using *DESeq2* (Love et al., 2014). Given the high degree of collinearity between covariates (Fig. 1a), and the lower sample size per group (N=4-5 per cell line per DHT status), we used a simplified model for differential expression analysis:

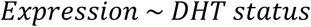

Features with a nominal P-value below 0.05 and an absolute fold change greater than 1.5 were retained for downstream analysis. Patterns across sex, ASD diagnosis, seizure status, 16p11.2 deletion status were assessed using a random effects model, offered through the *glmmSeq* package, where the cell line was set as the random intercept (Lewis et al., 2022):

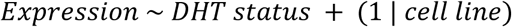

For this analysis, features with a nominal P-value below 0.05 were considered differentially expressed. A fold change filter was not applied given the random effects framework was applied to detect consistency in DHT effects rather than identifying large magnitude gene expression changes. Overlap between gene sets was determined using the Jaccard coefficient:

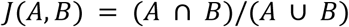

Where (𝐴 ∩ 𝐵) is the intersection of two DEG lists and (𝐴 ∪ 𝐵) represents the unique elements in 𝐴 and 𝐵.

### Functional enrichment analysis

Gene ontology (GO) enrichment analysis, using the *fenr* package, allowed use to inform biological interpretation of differentially expressed genes (Gierlinski, 2025). We examined associations with GO biological processes, cellular components, and molecular function terms. Enrichment terms were filtered to retain terms with a false-discovery-rate (FDR) adjusted p-value less than 0.1 and more than 2 genes that overlapped with the term geneset. We identified parent ontology terms using *rrvgo*, which groups functionally redundant GO terms based on their semantic similarity (Sayols, 2023). The semantic similarity matrix between enriched terms was calculated using the relevance information content method implemented through GOSemSim (Yu et al., 2010). Term significance was assigned using the −𝐿𝑜𝑔_10__(𝐹𝐷𝑅 − 𝑎𝑑𝑗𝑢𝑠𝑡𝑒𝑑 𝑝 − 𝑣𝑎𝑙𝑢𝑒), and redundant terms were collapsed with a similarity threshold of 0.7.

### Gene set profiling analysis

Geneset-level profiling was performed to characterize coordinated expression patterns across biological components, cell-type enriched genes, and biological processes. Cell component genesets were obtained using the *msigdbr* package, while cell type-enriched genes were sourced from the Human Protein Atlas (HPA) (Dolgalev, n.d.; *The Human Protein Atlas*, n.d.; Uhlén et al., 2015). For each geneset, we summarized expression using either the median expression (cell component and cell type-enriched genesets) or the first principal component (PC1) (for biological process genesets). Associations between geneset profiles and DHT status were evaluated using random effects models offered through the *lme4* package (Bates et al., 2015):

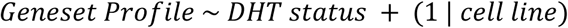

Associations below a nominal p-value threshold of 0.1 were considered significant. Moderators were incorporated into each model to test whether the associations depended on sex, ASD diagnosis, seizure status, or 16p11.2 deletion status:

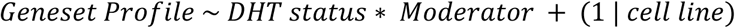

Interaction terms with a nominal p-value below 0.1 were considered significant. This framework enabled the evaluation of both main effects and moderated relationships.

## Data and code availability

All datasets and analysis code used in this study are available from the corresponding author upon reasonable request.

## Acknowledgments

Isogenic cell lines (F2-i and M2-i) were derived from cell line ID #17908 from the NIGMS Human Genetic Cell Repository at the Coriell Institute for Medical Research and provided within a cooperation by Prof. Benjamin Reubinoff (Hadassah Ein-Kerem Medical Center, Jerusalem, Israel). We express our appreciation for the opportunity to work with these cell lines.

We are grateful to all of the families at the participating Simons Searchlight sites as well as the Simons Searchlight Consortium, formerly the Simons VIP Consortium. We appreciate having access to phenotypic data on SFARI Base.

We wish to thank Enrique Ozcariz and Gabriel Alexander Vignolle from AIT Austrian Institute of Technology GmbH, Competence Unit Molecular Diagnostics, Austria for their support in analysing the DNA-methylation data. In addition, we thank Sabra Klein for her support in setting up the DHT treatment experiments.

RNA-seq was conducted at the Genetic Resources Core Facility (SCR_018669), Johns Hopkins Department of Genetic Medicine. We also thank George McNamara, PhD, the Ross Fluorescence Imaging Center (Johns Hopkins University), and the NIH shared instrumentation grant 1S10OD025244-01 for the use of the FV3000RS and the Johns Hopkins Multiphoton Microscopy Core for access to the LSM800.

This research was funded by the Deutsche Forschungsgemeinschaft (DFG, German Research Foundation) – Project number 507269789, the Wendy Klag Center at Johns Hopkins University and the Alternatives Research & Development Foundation (ARDF).

LS and AR were partially supported by NIH R01 R01ES034554. AR was also supported by the MSCRF TEDCO grant #2023-MSCRFD-6182, the Johns Hopkins MSTP 1T32GM136577-01, and the Johns Hopkins Environmental Health and Engineering 5T32ES007141-42.

## Author Contributions

Maren Schenke: Conceptualization, Methodology, Formal analysis, Investigation, Data Curation, Writing - Original Draft, Review & Editing, Visualization, Supervision, Project administration, Funding acquisition. Jason Laird: Methodology, Formal analysis, Investigation, Data Curation, Writing - Original Draft, Review & Editing, Visualization. Alex Rittenhouse: Methodology, Investigation, Writing - Review & Editing. Viktoriya Kucheryavenko: Investigation, Writing - Review & Editing. Winfried Neuhaus: Investigation, Writing - Review & Editing. Ou Chen: Investigation, Writing - Review & Editing. Sarven Sabunciyan: Resources, Writing - Review & Editing. Alexandra Maertens: Writing - Review & Editing, Supervision. Lena Smirnova: Methodology, Resources, Writing - Original Draft, Review & Editing, Supervision, Project administration, Funding acquisition

## Conflict of interest

LS is consulting for 28.bio (former Axosim), to whom the original bMPS model is licensed.

The remaining authors declare that the research was conducted in the absence of any commercial or financial relationships that could be construed as a potential conflict of interest.

