## Supplement for "Genetic Background and Sex Modulate Androgen Responses in Human Brain Microphysiological System"

### Supplementary information

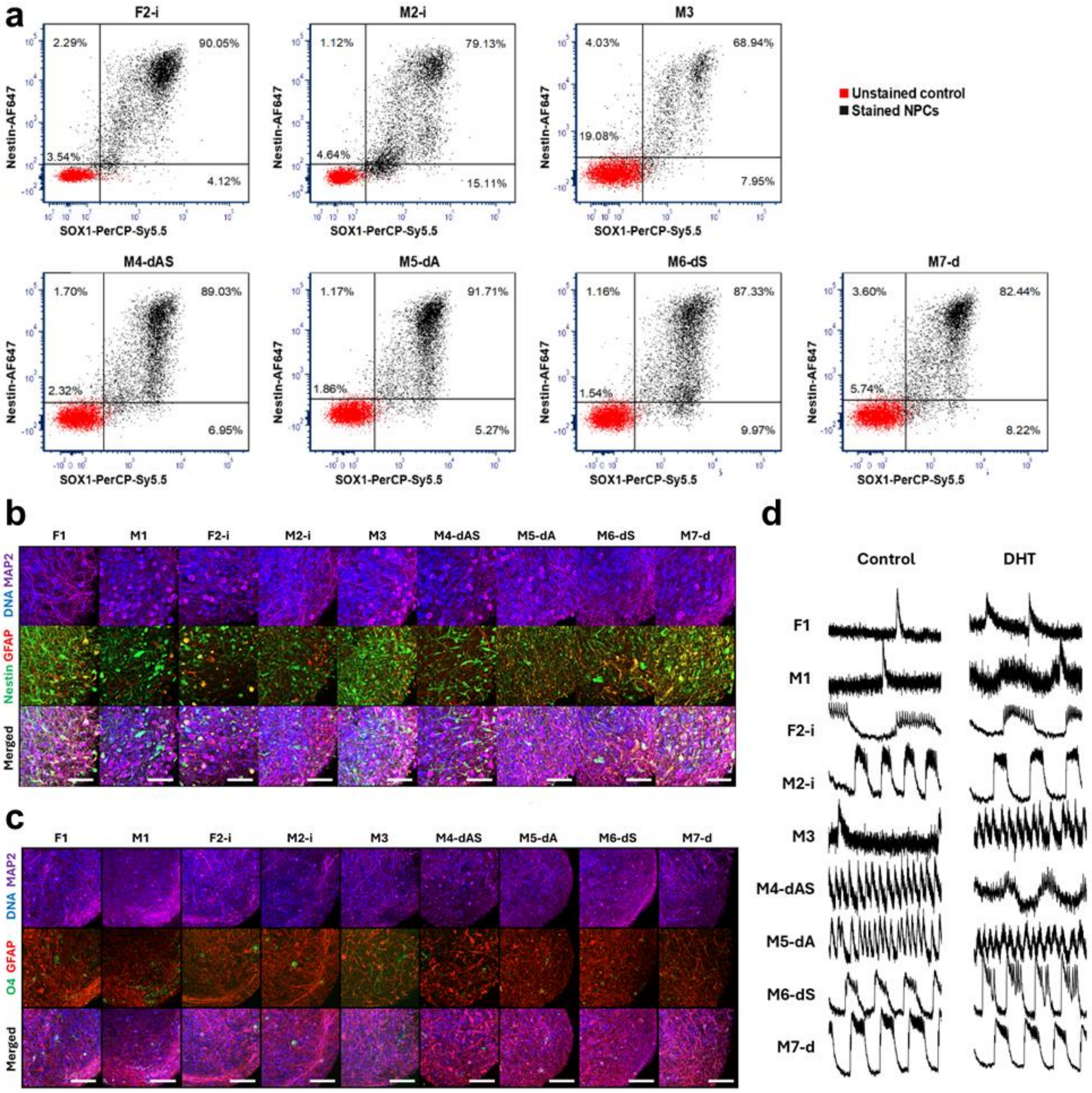


**Supplementary Fig. 1: Cell line and bMPS characterization. a)** Quantification via flow cytometry of SOX1 and Nestin positive NPCs generated from iPSC lines. **b)** Representative immunostaining of bMPS from all iPSC lines, at 4W of differentiation. Immature bMPS (4W) are positive for Nestin (neural stem cell marker), GFAP (glia marker), and MAP2 (neuronal marker). Scale bar is 50 µM **c)** Representative immunostaining of bMPS from all iPSC lines at 8W of differentiation. bMPS (8W) are positive for MAP2, GFAP and O4 (oligodendrocyte precursor cell marker). Scale bar is 100 µM. **d)** Representative plots of fluorescence over resting fluorescence (ΔF/F) determined via calcium imaging in bMPS differentiated for 8-9 weeks with 60 nM DHT or solvent treatment.

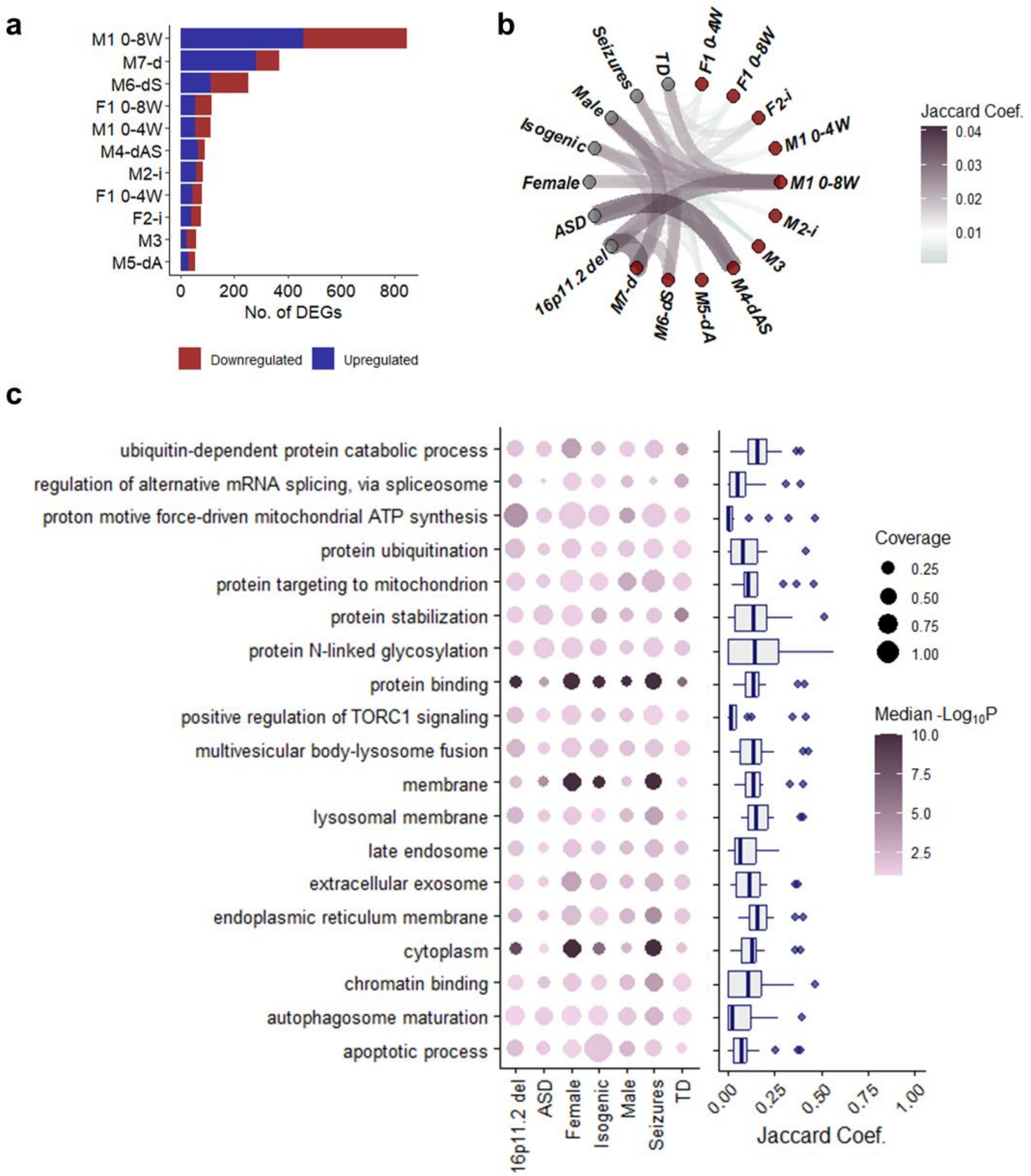


**Supplementary Fig. 2: Cell line transcriptional responses to DHT. a)** Number of DHT-induced DEGs ($P < 0.05$, $|FC| > 1.5$) across cell lines and exposure time points. If not explicitly labelled, cell lines were exposed to DHT for 8 weeks. **b)** Pairwise overlap between individual cell line DHT-associated DEGs and random effects DEGs colored by the Jaccard coefficient. **c)** Common DHT-associated GO enrichment terms (left) and the overlap in genes driving these common biological processes (right).

**Supplementary Table 1:** Donor characteristics and cell line attributes.

| Abbreviation | Source | Sex chromosomes | Genotype/ Phenotype | Age | Ancestry | Primary tissue and Reprogramming method |
|---|---|---|---|---|---|---|
| F1 | UK National Institute for Biological Standards and Control (NIBSC) | Female | Normal | Embryo | White | Lung epithelial, mRNA |
| M1 | Cedars-Sinai Medical Center Stem Cell repository | Male | Normal | 53 Y | White | PBMC, Episomal Plasmid |
| F2-i | Originally from Coriell cell repositories (GM17908), reprogrammed at Hadassah Stem Cell Research Center as described in Waldhorn et al. 2022 | Female | XX isolated from Patient with Klinefelter Syndrome with mosaic karyotype 47,XXY/46,XY/46,XX | 19 Y | Irish/English | B-Lymphocyte, nucleofected |
| M2-i | | Male | XY isolated from Patient with Klinefelter Syndrome with mosaic karyotype 47,XXY/46,XY/46,XX | | | |
| M3 | Cincinnati Children's Hospital | Male | Normal | 14 Y | Race: multiple, Ethnicity: Not Hispanic or Latino | PBMC,Sendai virus |
| M4-dAS | Simons Foundation Autism Research Initiative (SFARI) | Male | 16p11.2 Deletion ASD Seizures | 14 Y | | Fibroblasts, episomal |
| M5-dA | Simons Foundation Autism Research Initiative (SFARI) | Male | 16p11.2 Deletion ASD | 14 Y | | Fibroblasts, episomal |
| M6-dS | Simons Foundation Autism Research Initiative (SFARI) | Male | 16p11.2 Deletion Seizures | 5 Y | | Fibroblasts, episomal |
| M7-d | Simons Foundation Autism Research Initiative (SFARI) | Male | 16p11.2 Deletion | 8 Y | | Fibroblasts, episomal |

**Supplementary Table 2**: Antibodies used in this project

| Antibody | Host | Type | Manufacturer | Catalogue number | Dilution |
|---|---|---|---|---|---|
| GFAP | Rabbit | Polyclonal | Dako | #Z0334 | 1:400 |
| MAP2 | Chicken | Polyclonal | Invitrogen | #PA1-10005 | 1:5000 |
| O4 | Mouse | Monoclonal | R&D | #MAB1326 | 1:200 |
| Nestin | mouse | Monoclonal | Invitrogen | #14-9843-82 | 1:500 |
| AR | rabbit | recombinant | Proteintech | #81844-1-RR | 1:500 |
| AF647 anti Nestin | Mouse | Monoclonal | BD Bioscience | #560393 | 1:20 |
| PerCP-Cy anti SOX1 | Mouse | Monoclonal | BD Bioscience | #561549 | 1:20 |
| AF488 anti mouse IgG | Goat | Polyclonal | Invitrogen | #11001 | 1:500 |
| AF488 anti rabbit IgG | Goat | Polyclonal | Invitrogen | #A-11008 | 1:500 |
| AF647 anti chicken IgY | Goat | Polyclonal | Invitrogen | #A32933 | 1:500 |
| AF 568 anti rabbit IgG | Goat | Polyclonal | Invitrogen | #A-11011 | 1:500 |
| AF488 anti mouse IgM | Goat | Polyclonal | Invitrogen | #A-21042 | 1:500 |

**Supplementary Table 3**: Primer sequences used for RT-qPCR

| Gene name | Forward (5'-3') and reverse primer (5'-3') sequence | Efficiency [%] |
|---|---|---|
| **ACTB**<br>NM_001101.5 | CCTCGCCTTTGCCGATCC<br>CGCGGCGATATCATCATCCAT | 113 |
| **PPIA**<br>NM_021130.5 | CGCCGAGGAAAACCGTGTACT<br>CCTTGTCTGCAAACAGCTCAAAG | 94 |
| **AR**<br>NM_000044.6 | CTGTGCGCCAGCAGAAATGA<br>TTTCTTCAGCTTCCGGGCTCC | 107 |
| **ESR1**<br>NM_000125.4 | AGCACCCTGAAGTCTCTGGA<br>TGCTCCATGCCTTTGTTACTCA | 105 |
| **CYP19A1**<br>NM_001347252.2 | ACCCTTCTGCGTCGTGTCAT<br>ACCACGATAGCACTTTCGTCCAA | 101 |
| **SRD5A1**<br>NM_001047.4 | GGCTTGTGGTTAACGGGCAT<br>TTCAAATAAGCCTCCCCTTGGT | 102 |

**Supplementary Data 1**

DEGs for all cell lines following DHT treatment.

**Supplementary Data 2**

Random effects model results identifying genes with significant DHT effects across donors.

**Supplementary Data 3**

GO enrichment results from the random effects of DHT-associated genes, containing enriched processes per condition.

**Supplementary Data 4**

Baseline transcriptional differences between DHT-responders and non-responders.

**Supplementary Data 5**

GO enrichment analysis of baseline gene expression differences between DHT-responders and non-responders.

**Supplementary Data 6**

Differential expression results comparing sex chromosome effects (F2-i vs. M2-i) and DHT effects in the male isogenic line (M2-i v. DHT M2-i).

**Supplementary Data 7**

GO enrichment results for the sex chromosome vs. DHT comparisons.

**Supplementary Data 8**

DMPs following DHT treatment per cell line.

**Supplementary Data 9**

GO enrichment results of DMPs across cell lines.
